# Analysis of structure of the population, kinship coefficients and inbreeding trend depending on sex, type of breeding of Tatra Sheepherd dogs

**DOI:** 10.1101/2020.02.19.956045

**Authors:** Sweklej Edyta, Horoszewicz Elżbieta, Niedziółka Roman

## Abstract

The aim of the study was to analyse the structure of the population, kinship coefficients and inbreeding trend taking into account the sex, breeding system: champions (CH) and non-champions (nCH), breeding country: Poland (PL) and foreign country (Z) and the inbreeding degree of Tatra Shepherd dogs. Out of the currently registered 587 Tatra Shepherd dogs, 41.9% have been qualified for breeding. In the past decade, 1961 puppies were born, which corresponds to an average litter of 5.8 puppies. The breed’s inbreeding rate amounted to 6.34%, and for a 4-generation population was 6.68%. The highest inbreeding rate was found in nCH and PL groups consisting of both male and female dogs. The inbreeding rate was significantly higher in 2005-2014 compared to the years 1994-2004. The limit value F_X_ was exceeded for 25.65% of Shepherd dogs, and the critical value was exceeded for 11.52%. An increasing ancestor loss coefficient (AVK) was found, which may result in an increased number of inbred animals. In particular, it referred to female dogs in the nCH, PL, and F group, whereas a significant increase of AVK was observed in the group of male dogs from foreign kennels. The resulting COR values, respectively 55.58% for males and 55.44% for females, testify to insignificant inbreeding and suggest that breeders look for male inbreds. Studies have shown that there is no risk of inbred depression yet; however, the gene pool of the Tatra Shepherd dog breed has become noticeably restricted. In addition, leaving the stud book for the breed open must be considered due to an increase in the popularity of the breed, and thus an increase in mating.

## Introduction

There are about 500 million dogs around the world, including more than 63 million living in the EU. The highest number of dogs in Europe live in Russia – more than 15 million, in the Germany – 9,4 million, in the United Kingdom – 9 million and in the Poland 7,6 million. It is estimated that 25% of households in the European Union and 24% in the Europe keep at least one dog. In Poland and Romania, it is estimated that 42% of households have at least one dog, which puts Poland first in Europe. On the other hand, according to the American Veterinary Medical Association (AVMA), more than 70 million dogs live in the United States of America, while the American Pet Products Association (APPA) estimates their count as about 77.8 million. Both sources recount that smaller breeds are predominant in the USA. The World Canine Organization (Fédération Cynologique Internationale, FCI) classified 367 breeds, and Poland has currently 340 registered breeds split into hunting, guardian and shepherd dogs [1-5].

The oldest canine organization in Poland is the Polish Kennel Club (ZKwP – founded in 1938) which keeps Stud Books (KW) for Polish breeds and maintains the longest pedigree lines for all world recognized breeds, verifies pairs of parents and registered litter, conducts mental health tests and organizes renowned shows of purebred dogs. ZKwP is a patron of 5 national breeds including: Polish Grey Hound (FCI standard no. 333), Polish Hunting Dog (FCI standard no. 354), Polish Hound (FCI standard no. 52), Polish Lowland Sheepdog (FCI standard no. 251) and Tatra Shepherd Dog (FCI stanadard no. 252). In 1973 Fédération Cynologique Internationale (FCI), at the request of ZKwP, approved the Tatra Shepherd dog standard number 252 as a shepherd and guardian dog. In order to protect the exterior of dogs in Poland every year more than 120 purebred dog shows are organized, including 15 international ones with the participation of Tatra Shepherd dogs [6, 7]. Recently, dog shows have become increasingly popular, and the selection has been more oriented at the phenotype and grooming of dogs [8]. Studies show that such breeding leads to a loss of genetic variability in some breeds in Europe [9-14].

The Tatra Shepherd Dog breed has a number of relatives classified as separate breeds living along the Carpathians, Alps and Pyrenees. The territorial proximity between Tatra Shepherd dog and Slovak Cuvac suggest a close relationship of these two breeds. However, based on the cross-population studies, on the grounds of DNA polymorphism a genetic distance dendrogram was designed for six breeds of white shepherd dogs: Pyrenean Mountain Dog (France), Maremma Sheepdog (Italy), Akbash Dog (Turkey), Tatra Shepherd Dog (Poland), Slovak Cuvac (Slovakia), and the Kuvasz (Hungary). Studies have shown that Tatra Shepherd dogs are the closest relatives of Akbash dogs [7, 15, 16]. Despite their unique guarding efficiency, Tatra Shepherd Dogs indirectly contribute to green grazing of sheep and are a tourist amenity to the numerous visitors in the region of Podhale [1, 3, 17, 18]. The main features that predispose them to this type of work are endurance, strength, courage, distrust of strangers, resistance to various weather conditions, good hearing and sight, and a well-developed defensive instinct [19, 20].

Despite many genetically valuable Tatra Shepherd dogs having been taken abroad from Poland by their owners, the population seems to be reconstructed and quite even. However, it is still not sufficiently numerous, which prompts the monitoring of relationship in order to avoid inbreeding [21, 22].

Inbreeding increases the level of homozygotism of an animal, which has positive and less positive effects. Creating homozygous animals can reveal latent defects in recessive genes. Many authors claim that the loss of genetic variability and inbreeding contributes to the development of physical diseases, defects and disorders such as reduced fertility and prolificacy, the occurrence of lethal alleles in a litter, and lower offspring survival rate. In addition, anomalies may occur in the hip joint anatomy leading to dysplasia among house dogs, especially of large breeds [23-26]. The critical inbreeding rate is 12.5%. The inbreeding coefficients for a breed range from about 0.82% for Golden Retrievers, 2% for German Shepherd Dogs, 4,5% for Bullmastiff Dogs, through 9% for Great Danes to 26% for Nova Scotia Duck Tolling Retrievers and as much as 37% for Polish Hounds [27-33]. The F_x_ coefficient is supplemented by information about the content of a unique pool of genes revealed by the Ancestor Loss Coefficient (AVK). AVK makes it possible to assess inbreeding in previous generations and indicates a percentage ratio of unique ancestor to the total number of ancestors for a specific number of generations [9, 34-36].

The variations in the population of Tatra Shepherd Dogs have been shown by Radko et al. [37]. The DNA polymorphism analysis demonstrated a considerable genetic differentiation of microsatellite markers and the lack of inbreeding in the analysed population of Tatra Shepherd dogs. Other studies carried out in Poland [21, 38, 39] confirm the differentiated level of inbreeding with an upward trend, depending on the sex and age as well as the region of occurrence of the Tatra Shepherd Dog breed. It should be accepted that genetic equilibrium rarely occurs even in natural populations where natural choice and selection take place [40, 41].

The studies aimed at analyzing the structure of the population, estimating the relationship and inbreeding trend based on pedigrees taking into account features such as: sex, breeding type (champions, non-champions) and origin (domestic and foreign) of Tatra Shepherd dogs.

## Material and methods

### The origin of the Tatra Shepherd breed

The first pedigree Tatra Shepherds in Poland were born in 1957 in Łeba, in the kennel run by Danuta Hryniewicz. They were the descendants of dogs she had owned since 1935, registered with the Polish Association of Pedigree Dog Breeders. During the first International Pedigree Dog Show in Poland held in Poznań in 1962, only two Tatra Shepherds were shown, but in the following years the number of those dogs increased continuously. In 1973, the Fédération Cynologique Internationale (FCI), at the request of ZKwP (Polish Kennel Club) approved a breed standard of Tatra Shepherd dog under number 252. In 1979–1980 Tatra Shepherds were the most numerous Polish breed of dogs registerd with ZKwP (in total 329 dogs) [6,7]. After 1981 Clubs or sections associating Tatra Shepherd breeders were established abroad. The kennels comprised from several hundred (Netherlands, Belgium, Germany) to several dozen dogs (USA, France, Austria, Norway, Finland) [6, 42].

### Genetic analysis

Analysis of the indicators of genetic variation of the population covered dogs entered into the Stud Book (non-pedigree parents) and dogs with 1 to 18-generation pedigrees within the same breed. A pedigree database containing data of 505 Tatra Shepherds (202 male dogs and 303 female dogs) born in 1964-2014 was developed and used to identify a standardized population with full 4-generation pedigrees totalling 194 dogs, including 82 males and 112 females born in 1994-2014. The Wright coefficients of inbreeding (F_x_) and the ancestor loss coefficient (AVK) were estimated for all animals split according to: sex, having a champion title (CH) or not having a champion title (nCH), origin: from Polish kennels (POL) and from foreign kennels (Z). In addition, the coefficient of kinship between the proband and its direct primary ancestors (R_XA_) and secondary ancestors (R_XY_) was estimated.

### Statistical analysis

The inbreeding coefficient, the kinship coefficient of the proband and its direct primary ancestors (R_XA_) and secondary ancestors (R_XY_) is calculated according to the formulas [40, 43-45]:

Inbreeding coefficient:

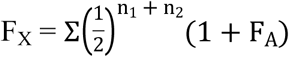

Secondary kinship:

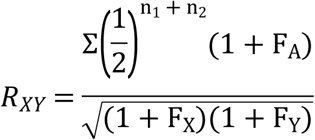

Primary kinship:

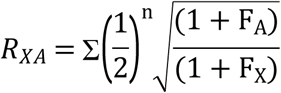

Key to formulas:

F_X_ – inbreeding coefficient of specimen _X_,

F_Y_ – inbreeding coefficient of specimen _Y_,

F_A_ – inbreeding coefficient of a common ancestor _A_

R_XA_ – coefficient of kinship between specimen _X_ and its ancestor _A_,

R_XY_ – coefficient of kinship between specimens _X_ and _Y_,

n_1_, n_2_ – number of paths from the parents of specimen X to an ancestor A shared by the parents.

The calculations were made using the Pedigree Explorer software (Wild Systems P/L, Australia). Secondary kinship was calculated with the support of the CFC programme [46]. The results of the study were subjected to an analysis of variance taking the following features into account: sex, champion (CH), non-champion nCH), domestic kennel (PL), and foreign kennel (Z). The data are presented as means (SE) with their standard deviations (SD). The significance of differences between the mean values for respective groups was verified by means of Tukey’s test (p≤0,001) using Statistica 12 software [47].

## Results

### Population of Tatra Shepherd dogs

The population of Tatra Shepherd dogs has been growing both in domestic and foreign kennels. Since 2003 a year-on-year increase has been observed in the number of Tatra Shepherd dogs newly registered with the divisions of the Polish Kennel Club. Every year, more females than male dogs are registered. At the end of 2011, the number of registered Tatra Shepherds was 473 (184 males and 289 females), and 112 Shepherds, including 46 male dogs and 66 female dogs, were qualified for breeding (Table 1). As at 31 December 2016, out of 587 registered dogs (221 males and 366 females) as many as 246 (83 males and 163 females) were qualified for breeding, which corresponded to 41.9% of all the registered animals of this breed. Over five years only (2011-2016), 1961 puppies were born from 337 litters, which accounts for an average of about 5.8 puppies per litter. The analysis of the trend until 2021 shows a constant growth in the breed population both among male dogs (+8.30%) and female dogs (+17.93%). The largest growth is observed among breeding dogs. This is a positive symptom of recognizing the Tatra Shepherd dog in the population. Among other things, it is due to its suitability as a shepherd and guard dog but also due to the decision of the authorities to open a stud book for the breed in order to extend the gene pool. In 2016 there were 121 registered Tatra Shepherd dog breeders in Poland. The highest number, that is, 40 are registered in the Lesser Poland voivodeship **–** in the region of Podhale, which corresponds to 33.06%.

**Table 1.** The population of registered Tatra Shepherd dogs within the time interval 2011-2016 (number of dogs) and the forecasted growth in 2017-2021.

| Year | Adult dogs |  |  |  | Puppies |  |
| --- | --- | --- | --- | --- | --- | --- |
|  | Dogs in total | Breeding Dogs | Bitches in total | Breeding Bitches | Dogs | Bitches |
| 2011 | 184 | 46 | 289 | 66 | 136 | 150 |
| 2012<sup>1</sup> | - | - | - | - | - | - |
| 2013 | 218 | 81 | 309 | 152 | 190 | 201 |
| 2014 | 222 | 85 | 308 | 150 | 193 | 195 |
| 2015 | 220 | 89 | 337 | 152 | 205 | 224 |
| 2016 | 221 | 83 | 366 | 163 | 259 | 208 |
| Trend until 2021<sup>2</sup> | +8.30 | +47.80 | +17.93 | +56.18 | +29.81 | +19.23 |

Data collected from all branches of the Cennel Club for the Tatra Shepherd dog [6], <sup>1</sup> No confirmed data, <sup>2</sup> Trend values expressed in %

### Pedigree analysis and occurrence of inbreeding

The studies have shown a considerable degree of inbreeding within the breed (Table 2). The average inbreeding rate F_X_ throughout the analysed population from the stud book (KW) and with 1 to 18-generation pedigrees was 2.93%. Only for inbred animals (F_X_) it amounted to 6.4%. For a standardized 4-generation population, the inbreeding rate was 6.68%, and 6.85% for inbred animals.

**Table. 2.** The average inbreeding rate of the Tatra shepherds population.

| Specification | Group | The whole population |  | 4 generational pedigree |  |
| --- | --- | --- | --- | --- | --- |
|  |  | N | mean± SD | N | mean± SD |
| **F<sub>x</sub>**<br>**Total** | **Dogs** | 198 | 3,14 ± 0.46 | 82 | 6,52 ± 0.48 |
|  | **Bitches** | 293 | 2,79 ± 0.42 | 112 | 6,79 ± 0.44 |
|  | **Total** | 491 | 2,93 ± 0.44 | 194 | 6,68 ± 0.45 |
| **F<sub>x</sub>**<br>**Inbred animals** | **Dogs** | 99 | 6,29 ± 0.47 | 81 | 6,60 ± 0.47 |
|  | **Bitches** | 128 | 6,38 ± 0.42 | 108 | 7,04 ± 0.42 |
|  | **Total** | 227 | 6,34 ± 0.44 | 189 | 6,85 ± 0.45 |

N – Number, SE – group average, SD - Standard deviation, F<sub>x</sub> – Inbreeding rate expressed in %

For the purposes of further calculations, the dogs with 4-generation pedigrees were divided into: champions (CH), non-champions (nCH) and dogs bred in Poland (PL) and abroad (Z) (Table 3). The average inbreeding value for Tatra shepherd dogs with full, 4-generation pedigrees, born in 1994-2014 amounted to 6.52% for males and 6.79% for females. The highest inbreeding rate was observed among males (7.08%) and females (7.12%) in the group of non-champions (nCH) and among males (6.75%) and females (6.91%) born in Polish kennels (PL). To get a full picture of changes in the inbreeding rate of this breed, two periods of about ten years each were selected: 1994-2004 and 2005-2014. The level of inbreeding among Tatra Shepherd dogs born in 1994-2004 was 5.87% for males and 4.88% for females, whereas in 2005-2014 it was 6.94% and 8.22%, respectively. Statistically significant differences were identified only for female dogs born in Poland, while they were highly significant for females born abroad and non-champion females, as well as all female dogs together. The significantly lowest (p≤0.001) inbreeding rate (388%) was recorded for female dogs born in 1994-2004 outside Poland in comparison to the group of female dogs born in 2005-2014. The highest inbreeding rate was found in the group of nCH females in 2005-2014 and was significantly (p≤0.001) higher in comparison to the nCH group in the years 1994-2004. A decreased rate of inbreeding among male dogs born in 2005-2014 in comparison to dogs born in 1994-2004 was observed only in the group of champion dogs. Other groups showed an upward but statistically insignificant trend.

**Table 3.** Values of the inbreeding rate of the standardized population of Tatra Shepherd dogs, taking into account their sex, group and years of study.

| Analyzed years | Dogs |  |  |  |  | Bitches |  |  |  |  |
| --- | --- | --- | --- | --- | --- | --- | --- | --- | --- | --- |
|  | Total | CH | nCH | PL | Z | Total | CH | nCH | PL | Z |
|  | N=82 | N=31 | N=51 | N=71 | N=11 | N=112 | N=28 | N=84 | N=96 | N=16 |
| **F<sub>x</sub>**<br>(1994-2004) | 5.87 | 6.04 | 6.62 | 6.22 | 4.33 | 4.88<sup>A</sup> | 5.38 | 4.60<sup>A</sup> | 5.08<sup>a</sup> | 3.88<sup>A</sup> |
| ±SD | 0.50 | 0.41 | 0.51 | 0.48 | 0.61 | 0.46 | 0.40 | 0.46 | 0.46 | 0.38 |
| **F<sub>x</sub>**<br>(2005-2014) | 6.94 | 4.90 | 7.58 | 7.06 | 5.86 | 8.22<sup>B</sup> | 6.46 | 8.58<sup>B</sup> | 8.19<sup>b</sup> | 8.40<sup>B</sup> |
| ±SD | 0.47 | 0.41 | 0.50 | 0.45 | 0.49 | 0.42 | 0.38 | 0.42 | 0.43 | 0.37 |
| **F<sub>x</sub>**<br>(1994-2014) | 6.52 | 5.60 | 7.08 | 6.75 | 5.02 | 6.79 | 5.80 | 7.12 | 6.91 | 6.14 |
| ±SD | 0.48 | 0.41 | 0.50 | 0.46 | 0.58 | 0.44 | 0.39 | 0.45 | 0.44 | 0.37 |

N – Number, SE – group average, ±SD - Standard deviation, F<sub>x</sub> – Inbreeding rate expressed in %, CH - Champion, nCH- Untitled champion, PL – Bred in Poland, Z –Bred abroad
<sup>a,b</sup> – Sex averages marked with different letters in columns differ significantly at p≤0,05
<sup>A,B</sup> – Sex averages marked with different letters in columns differ significantly at p≤0,001

The intensity of inbred classes (F_X_) among Tatra Shepherd dogs shows the lowest number of specimens in the class of non-related animals and in particular male dogs (Table 4). The limit value is 6.5% and the critical value 12.5%. The analysis shows that 25.65% of Tatra Shepherd dogs, including 23.75% males and 27.03% females exceeded the limit inbreeding value. On the other hand, 11.52% (respectively 12.50% males and 10.81% females) exceeded the critical inbreeding value.

**Table 4.** Intensity of inbred classes for Tatra Shepherd dogs.

| Inbreeding classes | Total | Dogs | Bitches |
| --- | --- | --- | --- |
| $F_x=0\%$ | 2.62 | 1.25 | 3.60 |
| $0 < F_x < 1,5\%$ | 5.24 | 11.25 | 0.90 |
| $1,5 < F_x < 6,5\%$ | 54.97 | 51.25 | 57.66 |
| $6,5\% < F_x < 12,5\%$ | 25.65 | 23.75 | 27.03 |
| $12,5 < F_x$ | 11.52 | 12.50 | 10.81 |

Data $F_x$ represent the percentage share in the group

The average ancestor loss coefficient (AVK) within the entire analysed population (Table 5) entered in the stud book (KW) and with 1 to 18-generation pedigrees amounted to 92.30%, including 84.81% for inbred animals. However, for a standardized 4-generation population it is 82.61%, including 82.31% for inbred animals.

**Table 5.** The average ancestor loss coefficient in the population of Tatra Shepherds dogs.

| Specyfication | Group | The wole population |  | 4 generational pedigree |  |
| --- | --- | --- | --- | --- | --- |
| | | N | mean ± SD | N | mean ± SD |
| AVK<br>Total | Dogs | 198 | 91.13 ± 9.88 | 82 | 82.93 ± 9.46 |
| | Bitches | 293 | 92.42 ± 9.85 | 112 | 82.38 ± 9.95 |
| | Total | 491 | 92.30 ± 9.85 | 194 | 82.61 ± 9.75 |
| AVK<br>Inbred animals | Dogs | 99 | 85.12 ± 9.44 | 81 | 82.76 ± 9.40 |
| | Bitches | 128 | 84.57 ± 9.64 | 108 | 81.98 ± 9.52 |
| | Total | 227 | 84.81 ± 9.53 | 189 | 82.31 ± 9.65 |

Data area mean in % ± SD, AVK – Ancestor loss coefficient

The values of the ancestor loss coefficient for Tatra Shepherd dogs with full, 4-generation pedigrees split into groups: CH, nCH, PL and Z are presented in Table 6. The highest AVK was noted for male dogs (84.30%) and female dogs (85.83%) from the CH group, and the lowest for male dogs from the Z group (81.39%) and female dogs from the nCH group (81.23%). A higher AVK means less inbreeding. The AVK coefficient increased significantly (p≤0.05) in the group of dogs from foreign kennels to 93.30% in 2005-2014. AVK values were significantly decreased over the analysed years in the groups: nCH, PL, Z for female dogs to the following level, respectively: 77.17% (p≤0.001), 78.96% (p≤0.05), 77.25% (p≤0.001). It can be concluded that the ancestor loss coefficient and the inbreeding rate were growing in the group of female dogs.

**Table 6.** Values of the ancestor loss coefficient of the standardized population of Tatra Shepherd dogs, taking into account their sex, group and years of study.

| Analyzed years | Dogs |  |  |  |  | Bitches |  |  |  |  |
| --- | --- | --- | --- | --- | --- | --- | --- | --- | --- | --- |
|  | Total | CH | nCH | PL | Z | Total | CH | nCH | PL | Z |
|  | N=82 | N=31 | N=51 | N=71 | N=11 | N=112 | N=28 | N=84 | N=96 | N=16 |
| AVK<br>(1994-2004) | 83.33 | 83.33 | 83.33 | 83.87 | 81.43<sup>a</sup> | 87.57<sup>A</sup> | 86.47 | 88.17<sup>A</sup> | 87.08<sup>a</sup> | 90.00<sup>A</sup> |
| ±SD | 9.54 | 9.12 | 9.76 | 9.45 | 9.70 | 9.89 | 8.81 | 11.02 | 10.21 | 11.34 |
| AVK<br>(2005-2014) | 82.67 | 85.63 | 81.67 | 82.81 | 93.30<sup>b</sup> | 78.49<sup>B</sup> | 84.85 | 77.17<sup>B</sup> | 78.96<sup>b</sup> | 77.25<sup>B</sup> |
| ±SD | 8.97 | 8.89 | 9.32 | 9.31 | 9.56 | 9.98 | 8.70 | 9.23 | 8.94 | 10.80 |
| AVK<br>(1994-2014) | 82.93 | 84.30 | 82.09 | 83.19 | 81.39 | 82.38 | 85.83 | 81.23 | 82.18 | 84.24 |
| ±SD | 9.46 | 9.09 | 9.59 | 9.40 | 9.67 | 9.95 | 8.71 | 10.08 | 9.81 | 11.02 |

Designations as in Table 3, AVK – Ancestor loss coefficient

The estimated kinship in the breed is presented by Table 7. It turned out that the average kinship between probands and their parents in the direct male line was 55.58%, and in the female line 55.44%. Values above 50% indicate that each line, both paternal and maternal, was characterised by a certain degree of inbreeding. A higher kinship coefficient for the paternal line may suggest that breeders look for male line inbreds.

**Table 7.**
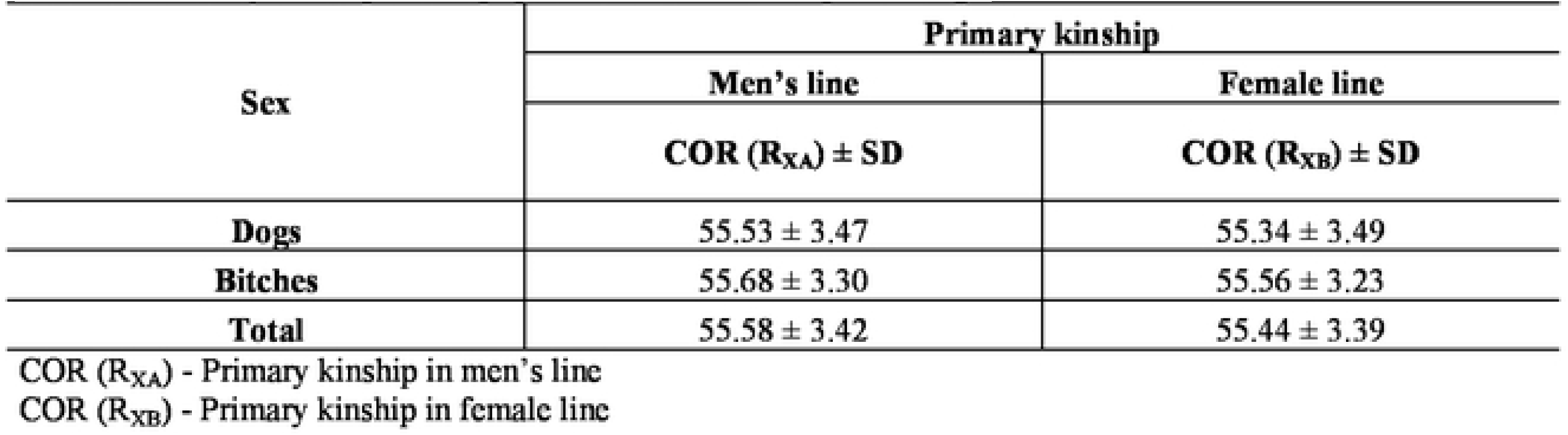
Primary kinship in the population of Tatra Shepherd dogs.

In the population of Tatra Shepherd dogs with 1 to 18-generation pedigrees (Table 8), an average secondary kinship between the proband and its parents was stated at the level of 7.9% for all mixed pairs and 12.15% for inbred pairs. Considering the secondary kinship for mixed pairs (male and female) of dogs with full, four-generation pedigrees, the following results were obtained: 12.72% for all the analysed pairs and 13.05% for inbred pairs.

**Table 8.**
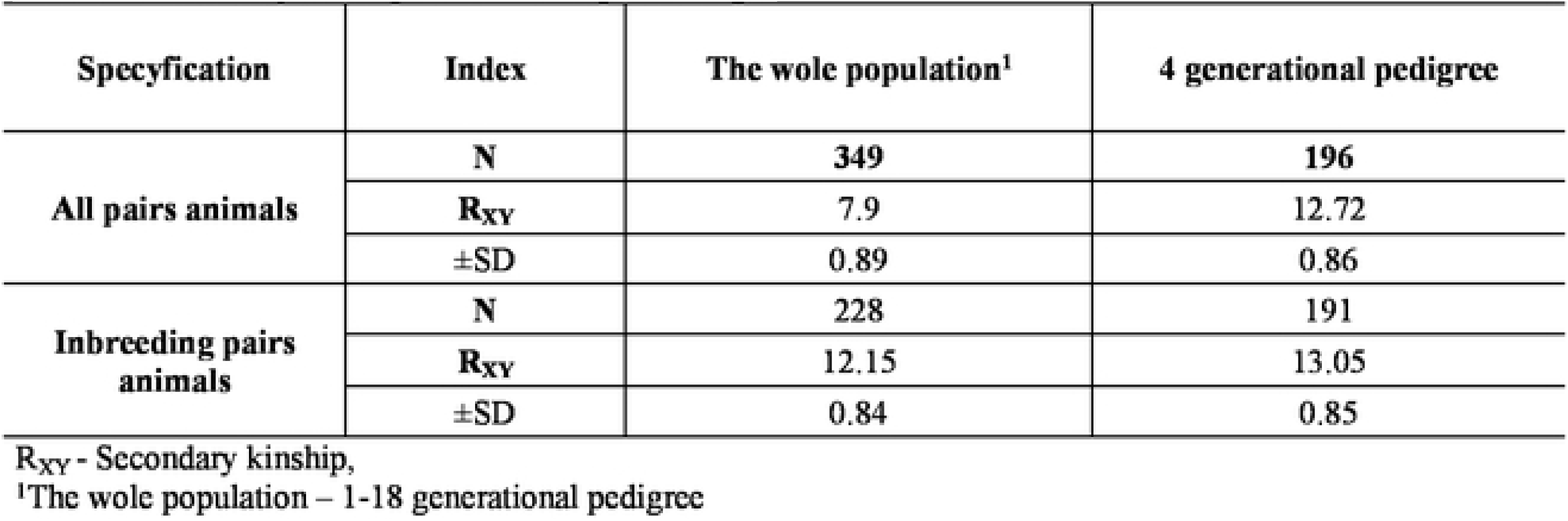
Secondary kinship of Tatra Shepherd dogs.

## Discussion

In 1999-2002 an average annual upward trend was noted among Tatra Shepherd dogs registered in Poland amounting to 7.4%, and in 2003 there was a downward trend of 36% in comparison to the previous year. In 2007, 21% of breeding female dogs were used for reproduction [48]. As a result of studies, it was established that in 2004-2014 the average annual trend among the registered Tatra Shepherds was upward in relation to the previous year by 11.9%, whereas in the discussed period both increases and decreases were observed. The largest increase was observed in 2004 in comparison to the previous year (69.2%), and the largest decrease (14.9%) in 2007. In 2011-2016, the kennels used on average 45% of breeding female dogs, and the highest percentage, i.e. 49%, was recorded in 2016.

The number of dogs shown at dog shows could be a measure of the breed’s popularity. The groups of breeds that are popular in Poland such as Retrievers (Golden or Labrador) or Yorkshire Terriers during the shows often include more than 100 animals [49]. Thus, the Tatra Shepherd dog cannot be classified as a popular breed, but it can be stated that the population of Tatra Shepherd dogs remained at a comparable level in 2005-2014, and the average number of dogs shown at dog shows was 21, with an average annual upward trend of 7.1%. Attendance at the most prestigious club dog shows was on average 56 dogs in 2005-2017, assuming the highest number, that is, 77 dogs in 2007 [6]. The number of Tatra Shepherd dogs abroad is relatively high (about 700 in the Netherlands, about 500 in Germany, and several dozen dogs in Belgium and Finland). The profit motive contributed to a high degree to the export of the best specimens, as puppies are often sold abroad at higher prices than in Poland [50-51].

Kalinowska et al. [21] demonstrated that half of the active population registered with the Krakow Division of ZKwP was inbred, whereas in a 4-generation population more than 23% of dogs were inbred (about 27% males and 20% females). The average inbreeding rate was 1.37%, and the rate estimated only among inbred animals is 5.85%. The highest inbreeding rate was 14.06%. The latest studies on the population consisting of 491 dogs showed that 46% of Tatra Shepherds (20% males and 26% females) were inbred, and in the standardized 4-generation population 40%, including 16% males and 22% females is of an origin connected with inbreeding. The average inbreeding rate was at the level of 2.93% for the entire population and among inbred animals it was 6.34%. The highest inbreeding rate was 20.31%. The studies by Kania-Gierdziewicz and Gierdziewicz [38] conducted for the Tatra Shepherd dog breed in Silesian voivodeship showed that 77.42% of the population is inbred (including 81.82% of males and 75% of females), and the average inbreeding rates (F_X_) are respectively 4.8% for all and 5.8% for inbred animals, whereas the average kinship coefficient is 11.5%. In another experiment carried out in 2015 on the population of Tatra Shepherd dogs living in the region of Podhale, Kania-Gierdziewicz et al. [22] found that the average inbreeding rate for the breed is 7.17%, and the average kinship coefficient is 18.2%. In addition, the inbreeding rate showed an upward trend. Similar F_X_ results at the level of 8.8% were obtained by Leroy et al. [12] for Pyrenean Shepherd and by Cecchi et al. [33] for the Bracco Italiano breed – F_x_ = 6,7%. A slightly higher inbreeding levels of the Boxer breed were above 10% and the inbreeding rate amounted to 0.14% per year [52]. In contrast Bullmastiff Dogs inbreeding coefficients ranged from 0 in 1980 to 0.054 in 1997, an overall increase in mean inbreeding coefficient is seen until the mid 1990’s, reaching 0.043 in 1995 and remain relatively stable to 0.044 in 2013 [13]. The current Norwegian Lundehund population is highly inbred and has lost 38.8% of the genetic diversity in the base population. The ancestor with the highest contribution in the pedigree is a female with 18 offspring born in the 1960s. Her contribution to the last cohort is 41%. Immediate actions are needed to increase the genetic diversity in the current Lundehund population. The only option to secure the conservation of this rare breed is to introduce individuals from foreign breeds as breeding candidates [11]. For Swedish protected dog breeds there is no correlation between average *F* and population size measured either as the size of the full pedigree or as the number of living dogs (coefficients of correlation, *r*, range from 0.00 to 0.53 and 0.15–0.60, respectively, with 0.07<P<1.00). Breeds like the Swedish lapphund and the Swedish vallhund, however, reach almost as high average *F-*0.09 -in spite of pedigree sizes of several thousand individuals, census sizes of well over 1000, and over 50 founders. Similarly, the Swedish elkhound and the Drever have pedigrees comprising over 50,000 dogs, with over 10,000 defined as alive in 2012 but average *F* is over 0.07 [10]. A study of genealogical parameters for a number of breeds in Australia found that the mean inbreeding coefficient ranged from 0 to 0.101 across 32 analysed breeds [53].

The AVK of the analysed Tatra Shepherds was satisfactory and it was above 91% for the general population of dogs and 84% for the inbred population. The period of analysis significantly contributed to a reduction in AVK, mostly for female dogs from groups nCH, PL, Z born in 2005-2014. Similar results for AVK were obtained for Newfoundland dogs – above 85% [35].

Secondary kinship R_XY_ within the entire population was lower than for inbred pairs by 4.25%. On the other hand, similar results were obtained for pairs with 4-generation pedigrees. Kalinowska et al. [21] obtained similar R_XY_ values for inbred pairs within the range 14.51-14.92% and lower for the entire population, 4.53-6.12%. Slightly lower R_XY_ results, around 3.91%, were estimated for the entire population of German Shepherd dogs from the region of Krakow [54].

With regard to quite a high level of inbreeding in the breed, new blood is recommended in breeding Tatra Shepherd dogs. The source of enriching the gene pool can be dogs entered in the stud book. Insofar as excessive inbreeding can contribute to inbreeding depression, the dogs entered in the stud book, despite their phenotype complying with the breed standard, are a mystery in terms of origin, genotype, possible inbreeding or why their ancestors were excluded from breeding. Thus, dogs to be entered into the stud book should be thoroughly selected and these should only be dogs having unquestionable typical physical and mental traits of the breed. Among all stud dogs in Poland mentioned in the Newsletter of the Tatra Shepherd Dogs Club, 61.11% did not leave a male line continuer [6, 21]. A way of enriching the line can be reaching the descendants of such dogs that are not shown at dog shows and obtaining genetic material by entering them into the stud book and then using them for breeding. Therefore, the import of Tatra Shepherd dogs bred in the Netherlands or France that show a low degree of kinship with dogs in Poland can be an excellent choice to abandon extensive inbreeding, and aim at improving the breed [22, 36, 37, 49].

This way of reducing the inbreeding is confirmed by studies concerning dog breeding and endangered breeds with small populations [13, 16, 33, 55].These observations are alarming since reduced genetic variation and inbreeding are generally associated with the loss of the adaptive potential and reduced options of effective selection [10, 52, 56]. We note that this rapid genetic diversity loss is parallel to an increasing requirement of dogs for a number of different purposes in the modern society [8, 57-59].

## Conclusion

In Poland, there are about 121 registered breeders of Tatra Shepeherd dogs, most of them in Lesser Poland voivodeship, i.e. 33.06% on a national scale. In 2016 a record-breaking number of puppies of this breed were born (467), while for the first time in the second decade of the 21^st^ century the number of male dogs born was 20% higher than the number of females.

Taking into account the inbreeding rate in the breed, it was concluded that no risk of inbreeding depression occurs yet but there is a trend to reduce the pool of genes. Therefore, it is suggested that the stud book for the Tatra Shepherd dog is left open because the breed has gained popularity year on year, so the number of matings will increase. Based on the results, it must be assumed that the maximum inbreeding rate (F_X_) in the analysed breed should not exceed 6.85%, whereas the ancestor loss coefficient (AVK) should not be lower than 76%.

## Author Contributions

**Conceptualization:** Edyta Sweklej, Elżbieta Horoszewicz, Roman Niedziółka.

**Data curation:** Edyta Sweklej, Elżbieta Horoszewicz.

**Formal analysis:** Edyta Sweklej, Elzbieta Horoszewicz, Roman Niedziółka.

**Funding acquisition**: Elżbieta Horoszewicz, Roman Niedziółka.

**Investigation**: Edyta Sweklej.

**Methodology**: Edyta Sweklej, Elżbieta Horoszewicz.

**Software**: Edyta Sweklej, Roman Niedziółka.

**Supervision**: Roman Niedziółka.

**Validation**: Edyta Sweklej, Elżbieta Horoszewicz.

**Visualization**: Elżbieta Horoszewicz, Roman Niedziółka.

**Writing – original draft**: Edyta Sweklej, Elżbieta Horoszewicz, Roman Niedziółka.

**Writing – review & editing**: Elżbieta Horoszewicz, Roman Niedziółka.

